# Evolution of research topics and paradigms in plant sciences

**DOI:** 10.1101/2023.10.02.560457

**Authors:** Shin-Han Shiu, Melissa D. Lehti-Shiu

## Abstract

Scientific advances due to conceptual or technological innovations can be revealed by examining how research topics have evolved. But such topical evolution is difficult to uncover and quantify because of the large body of literature and the needs of expert knowledge from a wide range of areas in any field. Here we used machine learning and language models to classify plant science citations into topics representing interconnected, evolving subfields. The changes in prevalence of topical records over the last 50 years reflect major research paradigm shifts and recent radiation of new topics, as well as turnovers of model species and vastly different plant science research trajectories among countries. Our approaches readily summarize the topical diversity and evolution of a scientific field with hundreds of thousands of relevant papers, and they can be applied broadly to other fields.

**Significance statement:** Changes in scientific paradigms are foundational for the advancement of science, but such changes are difficult to summarize, quantify, and illustrate. These challenges are exacerbated by the rapid, exponential growth of literature. Applying a combination of machine learning and language modeling to hundreds of thousands of published abstracts, we demonstrate that a scientific field (i.e., plant science) can be summarized as interconnected subfields evolving from one another. We also reveal insights into major research trends and the rise and decline in the use of model organisms in different countries. Our study demonstrates how artificial intelligence and language models can be broadly applied to understand scientific advances that inform science policy and funding decisions.

## Introduction

The explosive growth of scientific data in recent years is accompanied by a rapidly increasing volume of literature. These records represent a major component of our scientific knowledge and embody the history of conceptual and technological advances in various fields over time. Our ability to wade through these records is important for identifying relevant literature for specific topics, a crucial practice of any scientific pursuit (*1*). Classifying the large body of literature into topics can provide a useful means to identify relevant literature. In addition, these topics offer an opportunity to assess how scientific fields have evolved and when major paradigm shifts in took place. However, such classification is challenging because the relevant articles in any topic or domain can number in the hundreds or tens of thousands, and the literature is in the form of natural language, which takes substantial effort and expertise to process (*2, 3*).

In the last several years, there has been a quantum leap in natural language processing approaches due to the feasibility of building complex deep learning models with highly flexible architectures (*4, 5*). The development of large language models such as Bidirectional Encoder Representations from Transformers (BERT, (*6*)) and Chat Generative Pre-trained Transformer (ChatGPT, (*7*)) has enabled the analysis, generation, and modeling of natural language texts in a wide range of applications. The success of these applications is, in large part, due to the feasibility of considering how the same words are used in different contexts when modeling natural language (*6*). One such application is topic modeling, the practice of establishing statistical models of semantic structures underlying a document collection. Topic modeling has been proposed for identifying scientific hot topics over time (*1*), and it has also been applied to, for example, automatically identify topical scenes in images (*8*) and social network topics (*9*), and classify and extract features from biological data (*10*), and capture “chromatin topics” that define cell-type differences (*11*). Here we use topic modeling to ask how research topics in a scientific field have evolved and what major changes in the research paradigm have taken place, using plant science as an example.

## RESULTS

### Plant science corpora allow classification of major research topics

Plant science, broadly defined, is the study of photosynthetic species, their interactions with biotic/abiotic environments, and their applications. For modeling plant science topical evolution, we identified a collection of plant science documents (i.e., corpus) using text classification (see **Methods**). The best text classification model performed well (F1=0.961, F1 of a naïve model=0.5, perfect model=1, **fig. S1A,B**) and predicted 421,658 plant science citations, hereafter referred to as “plant science records” (**fig. S1C**, **Data S1**). Plant science records as well as PubMed articles grew exponentially from 1950 to 2020 (**Fig. 1A**), highlighting the challenges of digesting the rapidly expanding literature. We used the plant science records, to perform topic modeling, which consisted of four steps: representing each record as a BERT embedding, reducing dimensionality, clustering, and identifying the top terms by calculating class (i.e., topic)-based Term frequency-Inverse document frequency (c-Tf-Idf (*12*)). The c-Tf-Idf represents the frequency of a term in the context of how rare the term is to reduce the influence of common words. SciBERT (*13*) was the best model among those tested (**Data S2**) and was used for building the final topic model, which classified 372,430 (88.3%) records into 90 topics defined by distinct combinations of terms (**Data S3**). The topics contained 620 to 16,183 records and were named after the top 4–5 terms defining the topical areas (**Fig 1B**, **Data S3**). For example, the top five terms representing the largest topic, topic 61 (16,183 records), are “qtl,” “resistance,” “wheat,” “markers,” and “traits,” which represent crop improvement studies using quantitative genetics.

**Fig. 1.**
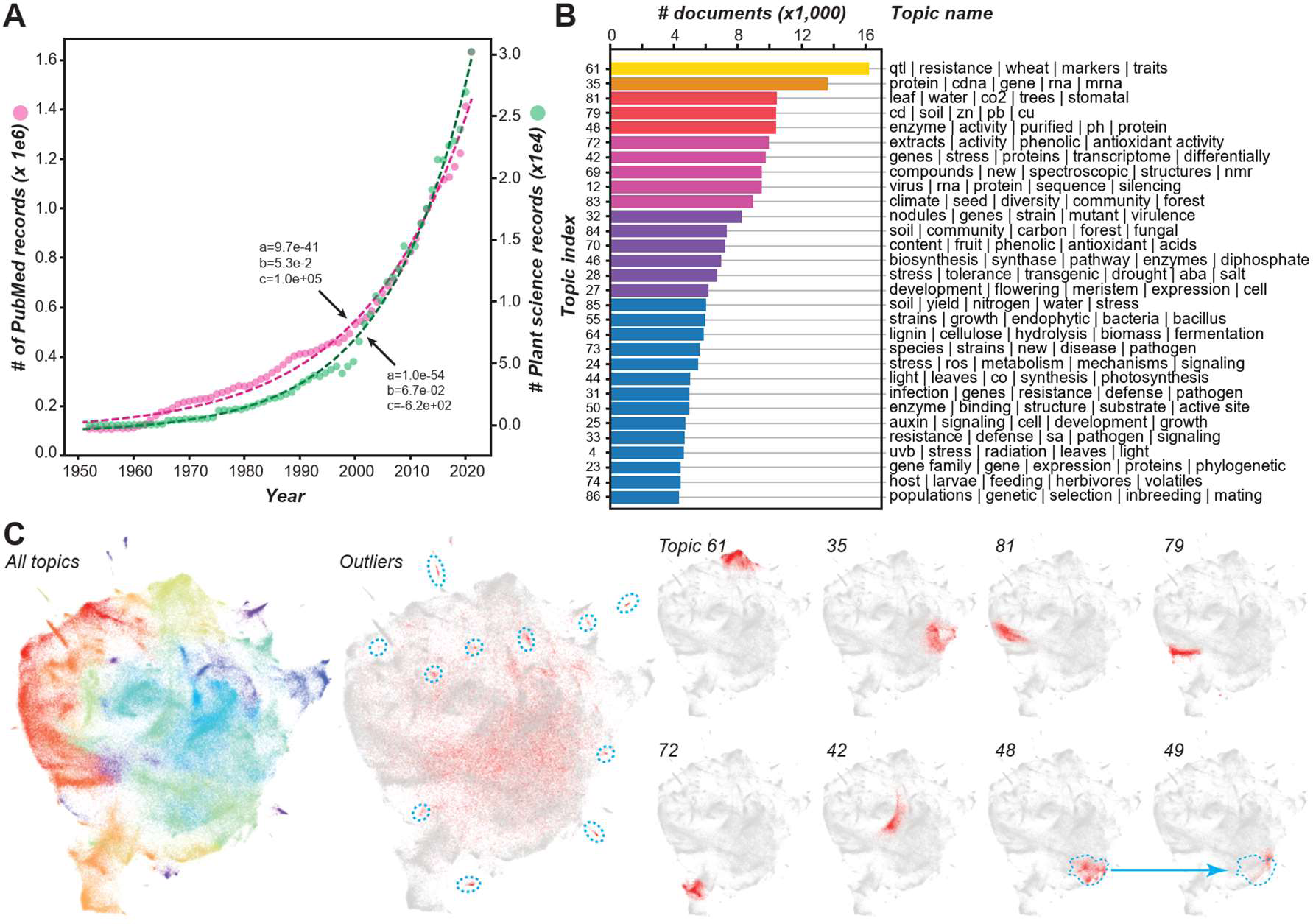
Change in the number of plant science records and research topics over time. **(A)** Numbers of PubMed (magenta) and plant science (green) records between 1950 and 2020. a, b, c: coefficients of exponential function, *y = ae^bx^*. **(B)** Numbers of documents for the top 30 plant science topics. Each topic is designated by an index number (left) and the top 4–6 terms with the highest cTf-Idf values (right). **(C)** Two-dimensional representation of the relationships between plant science records generated by Uniform Manifold Approximation and Projection (UMAP, (*14*)) using SciBERT embeddings of plant science records. All topics panel: different topics are assigned different colors. Outlier panel: UMAP representation of all records (gray) with outlier records in red. Blue dotted circles: areas with relatively high densities indicating topics that are below the threshold for inclusion in a topic. In the eight UMAP representations on the right, records for example topics are in red and the remaining records in gray. Blue dotted circles indicate the relative position of topic 48.

Records with assigned topics clustered into distinct areas in a two-dimensional (2D) space (**Fig. 1C**, for all topics, see **Data S4**). The remaining 49,228 outlier records not assigned to any topic (11.7%, middle panel, **Fig. 1C**) have three potential sources. First, some outliers likely belong to unique topics but have fewer records than the threshold (>500, blue dotted circles, **Fig. 1C**). Second, some of the many outliers dispersed within the 2D space (**Fig. 1C**) were not assigned to any single topic because they had relatively high prediction scores for multiple topics (**fig. S2**). These likely represent studies across sub-disciplines in plant science. Third, some outliers are likely interdisciplinary studies between plant science and other domains, such as chemistry, mathematics, and physics. Such connections can only be revealed if records from other domains are included in the analyses.

### Topical clusters reveal closely related topics but with distinct key term usage

Related topics tend to be located close together in the 2D representation (e.g., topics 48 and 49, **Fig. 1C**). We further assessed inter-topical relationships by determining the cosine similarities between topics using cTf-Idfs (**Fig. 2A**, **fig. S3**). In this topic network, some topics are closely related and form topic clusters. For example, topics 25, 26, and 27 collectively represent a more general topic related to the field of plant development (cluster *a*, **Fig 2A**). Other topic clusters represent studies of stress, ion transport, and heavy metals (*b*); photosynthesis, water, and UV-B (*c*); population and community biology (d); genomics, genetic mapping, and phylogenetics (*e*); and enzyme biochemistry (*f*, **Fig. 2A**).

**Fig. 2.**
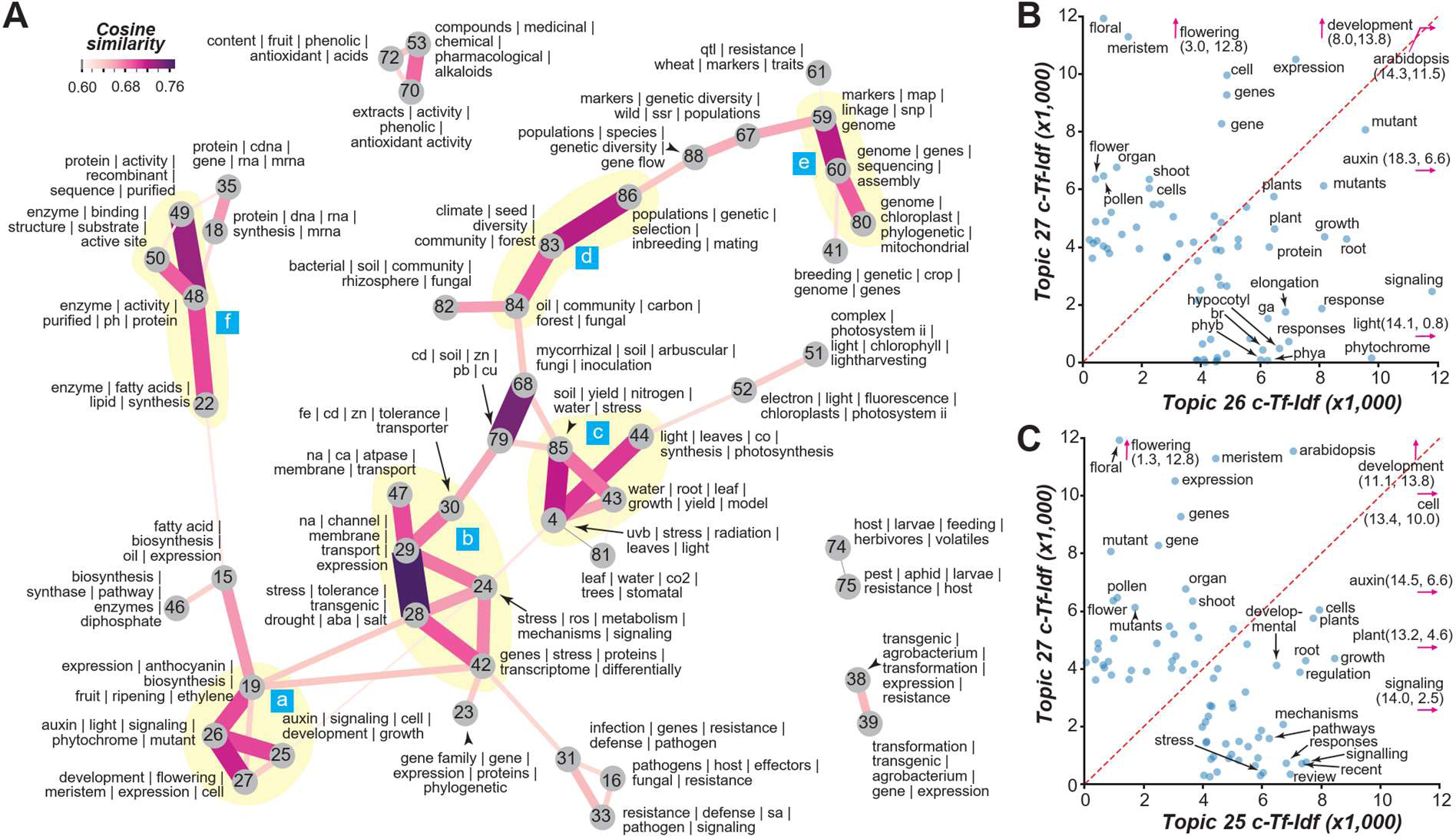
Inter-topical relationships. **(A)** Graph depicting the degrees of similarity (edges) between topics (nodes). Between each topic pair, a cosine similarity value was calculated using the cTf-Idf values of all terms. A threshold similarity of 0.6 was applied to illustrate the most related topics. For the full matrix presented as a heatmap, see **fig. S4**. The nodes are labeled with topic index numbers and the top 4–6 terms. The colors and width of the edges are defined based on cosine similarity. Example topic clusters are highlighted in yellow and labeled a through f (blue boxes). **(B, C)** Relationships between the cTf-Idf values of the top terms for topics 26 and 27 **(B)** and for topics 25 and 27. Only terms with cTf-Idf ≥ 0.6 are labeled. Terms with cTf-Idf values beyond the x and y axis limit are indicated by pink arrows and cTf-Idf values.

Topics differed in how well they were connected to each other, reflecting how general the research interests or needs are (see **Methods**). For example, topic 24 (stress mechanisms) is the most well connected with median cosine similarity=0.36, potentially because researchers in many sub-fields consider aspects of plant stress even though it is not the focus. The least connected topics include topic 21 (clock biology, 0.12) which is surprising because of the importance of clocks in essentially all aspects of plant biology (*15*). This may be attributed, in part, to the relatively recent attention in this area.

Examining topical relationships and the cTf-Idf values of terms also revealed how related topics differ. For example, topic 26 is closely related to topics 27 and 25 (**Fig. 2A**). Topics 26 and 27 both contain records of developmental process studies mainly in Arabidopsis (**Fig. 2B**); however, topic 26 is focused on the impact of light, photoreceptors, and hormones such as gibberellic acids (ga) and brassinosteroids (br), whereas topic 27 is focused on flowering and floral development. Topic 25 is also focused on plant development but differs from topic 27 because it contains records of studies mainly focusing on signaling and auxin with less emphasis on Arabidopsis (**Fig. 2C**). The similarities in cTf-Idfs between topics were also useful for measuring the editorial scope (i.e., diverse or narrow) of journals publishing plant science papers using a relative topic diversity measure (see **Methods**). For example, *Proceedings of the National Academy of Sciences, USA* has the highest diversity while *Theoretical and Applied Genetics* has the lowest (**fig. S4**). One surprise is the relatively low diversity of *American Journal of Botany*, which focuses on plant ecology, systematics, development, and genetics. The low diversity is likely due to the relatively larger number of cellular and molecular science records in PubMed, consistent with the identification of relatively few topical areas relevant to studies at the organismal, population, community, and ecosystem levels.

### Investigation of the prevalence of topics over time reveals topical succession

We next asked whether relationships between topics reflect chronological progression of certain subfields. To address this, we assessed how prevalent topics were over time using dynamic topic modeling (*16*). Because the number of plant science records has grown exponentially (**Fig. 1A**), the records were divided into 50 chronological bins each with ∼8,400 records to make cross-bin comparisons feasible (**Data S5**). Examining the prevalence of topics across bins revealed a clear pattern of topic succession over time (one topic evolved into another) and the presence of five topical categories (**Fig. 3**). The first is a stable category with six topics mostly established before the 1980s that have since remained relatively stable in terms of prevalence in the plant science records (**Fig. 3A**). These topics represent long-standing plant science research foci including studies of plant physiology (topics 4, 58, and 81), genetics (topic 61), and medicinal plants (topic 53). The second category contains eight topics established before the 80s that have mostly decreased in prevalence since (the early category, **Fig. 3B**). Two examples are physiological and morphological studies of hormone action (topic 45) and the characterization of protein, DNA, and RNA (topic 18). Unlike other early topics, topic 78 (paleobotany and plant evolution studies) experienced a resurgence in the early 2000s due to the development of new approaches and databases, and changes in research foci (*17*).

**Fig. 3.**
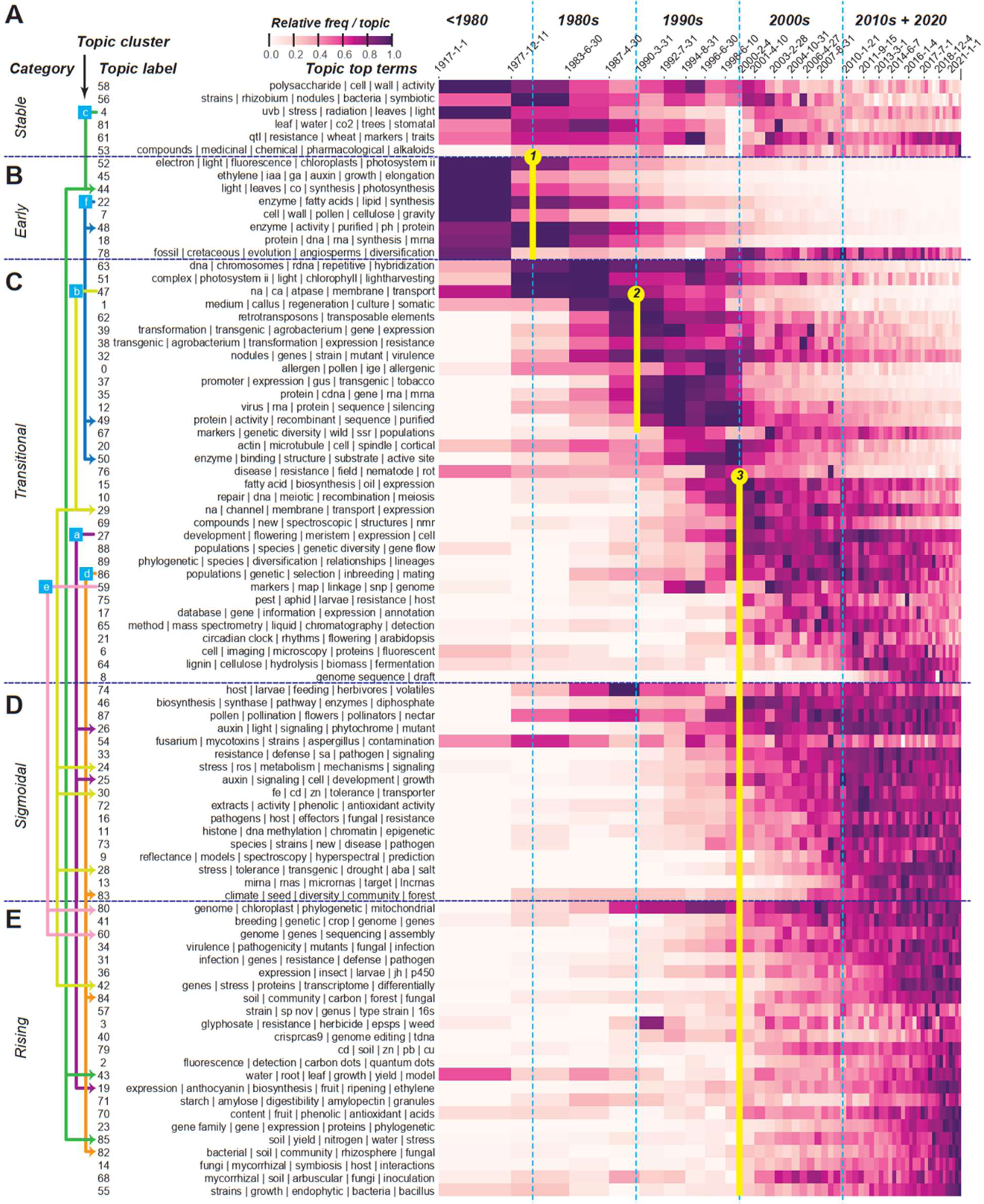
Topic evolution. **(A-E)** A heat map of relative topic frequency over time reveals five topical categories: **(A)** stable, **(B)** early, **(C)** transitional, **(D)** sigmoidal, and **(E)** rising. The x-axis denotes different time bins with each bin containing a similar number of documents. The sizes of all bins except the first are drawn to-scale based on the beginning and end dates. The y-axis lists different topics denoted by the label and top four to five terms. In each cell, the prevalence of a topic in a time bin is colored according to the min-max normalized cTf-Idf values for that topic. Light blue dotted lines delineate different decades. The arrows left of a subset of topic labels indicate example relationships between topics in topic clusters. Blue boxes with labels *a–f* indicate topic clusters, which are the same as those in **Fig. 2**. Connecting lines indicate successional trends. Yellow circles/lines 1*–*3: three major transition patterns.

The 33 topics in the third, transitional category became prominent in the 80s, 90s, or even 2000s, but have clearly decreased in prevalence (**Fig. 3C**). In some cases, the early and the transitional topics became less prevalent because of topical succession—re-focusing of earlier topics led to newer ones that either show no clear sign of decline (the sigmoidal category, **Fig. 3D**) or continue to increase in prevalence (the rising category, **Fig. 3E**). Consistent with the notion of topical succession, topics within each topic cluster (**Fig. 2**) were found across topic categories and/or were prominent at different time periods (indicated by colored lines linking topics, **Fig. 3**). One example is topics in topic cluster *b* (compare **Fig. 2, 3**); the study of cation transport (topic 47, transitional), prominent in the 80s and early 90s, is related to another transitional topic 29 (cation channels and their expression) peaking in the 2000s and early 2010s, sigmoidal topics 24 and 28 (stress response, tolerance mechanisms) and 30 (heavy metal transport), which rose to prominence in mid-2000s, and the rising topic 42 (stress transcriptomic studies), which increased in prevalence in the mid-2010s.

The rise and fall of topics can be due to a combination of technological or conceptual breakthroughs, maturity of the field, funding constraints, or publicity. The study of transposable elements (topic 62) illustrates the effect of publicity; the rise in this field coincided with Barbara McClintock’s 1983 Nobel Prize, but not with the publication of her studies in the 50s (*18*). The reduced prevalence in early 2000 likely occurred in part because analysis of transposons became a central component of genome sequencing and annotation studies, rather than dedicated studies.

### Three major topical transition patterns signify paradigm shifts

Beyond the succession of specific topics, three major transitions in the dynamic topic graph should be emphasized: (1) the decline of early topics in the late 70s and early 80s, (2) the rise of transitional topics in late 80s, and (3) the decline of transitional topics in the late 90s and early 2000s, which coincided with a radiation of sigmoidal and rising topics (yellow circles, **Fig. 3**). The large numbers of topics involved in these transitions suggest major paradigm shifts in plant science research. In transition 1, early topics decreased in prevalence in the late 70s to early 80s, which coincided with the rise of transitional topics over the following decades (circle 1, **Fig. 3**). For example, there was a shift from the study of purified proteins such as enzymes (early topic 48, **fig. S5A**) to molecular genetic dissection of genes, proteins, and RNA (transitional topic 35, **fig. S5B**) enabled by the wider adoption of recombinant DNA and molecular cloning technologies in late 70s (*19*). Transition 2 (circle 2, **Fig. 3**) can be explained by the following breakthroughs in the late 80s: better approaches to create transgenic plants and insertional mutants (*20*), more efficient creation of mutant plant libraries through chemical mutagenesis (e.g., (*21*)), and availability of gene reporter systems such as β-glucuronidase (*22*). Because of these breakthroughs, molecular genetics studies shifted away from understanding the basic machinery to understanding the molecular underpinnings of specific processes, such as molecular mechanisms of flower and meristem development and the action of hormones such as auxin (topic 27, **fig. S5C**); this type of research was discussed as a future trend in 1988 (*23*) and remains prevalent to this date. Another example is gene silencing (topic 12), which became a focal area of study along with the widespread use of transgenic plants (*24*).

Transition 3 is the most drastic: a large number of transitional, sigmoidal, and rising topics become prevalent nearly simultaneously at the turn of the century (circle 3, **Fig. 3**). This period also coincides with a rapid increase in plant science citations (**Fig. 1A**). The most notable breakthroughs included the availability of the first plant genome in 2000 (*25*), increasing ease and reduced cost of high-throughput sequencing (*26*), development of new mass spectrometry-based platforms for analyzing proteins (*27*), and advancements in microscopic and optimal imaging approaches (*28*). Advances in genomics and omics technology also led to an increase in stress transcriptomics studies (42, **fig. S5D**) as well as studies in many other topics such as epigenetics (topic 11), non-coding RNA analysis (13), genomics and phylogenetics (80), breeding (41), genome sequencing and assembly (60), gene family analysis (23), and metagenomics (82 and 55).

In addition to the three major transitions across all topics, there were also transitions within topics revealed by examining the top terms for different time bins (heatmaps, **fig. S5**). Taken together, these observations demonstrate that knowledge about topical evolution can be readily revealed through topic modeling. Such knowledge is typically only available to experts in specific areas and is difficult to summarize manually, as no researcher has a command of the entire plant science literature.

### Analysis of taxa studied reveals changes in research trends

Changes in research trends can also be illustrated by examining changes in the taxa being studied over time (**Data S6**). There is a strong bias in the taxa studied, with the record dominated by research models and economically important taxa (**fig. S6**). Flowering plants (Magnoliopsida) are found in 93% of records (**fig. S6A**), and the mustard family Brassicaceae dominates at the family level (**fig. S6B**) because the genus Arabidopsis contributes to 13% of plant science records (**Fig. 4A**). When examining the prevalence of taxa being studied over time, clear patterns of turnover emerged similar to topical succession (**Fig. 4B**, **fig. S6C,D, Methods**). For example, the increase in the number of Arabidopsis records coincided with advocacy of its use as a model system in the late 80s (*29*). While it remains a major plant model, there has been a decline in overall Arabidopsis publications compared to all other plant science publications since 2011 (blue line, **Fig. 4C**). Because the same chronological bins each with same numbers of records from the topic-over-time analysis (**Fig. 3**) were used, the decline here did not mean that there were fewer Arabidopsis publications—in fact the number of Arabidopsis paper remain steady since 2011. This decline means that Arabidopsis related publications were a relatively smaller proportion of plant science records. Interestingly, this decline took place much earlier (∼2005) and was steeper in the US, despite investment in the Arabidopsis 2010 Project by the US National Science Foundation (red line, **Fig. 4C**).

**Fig. 4.**
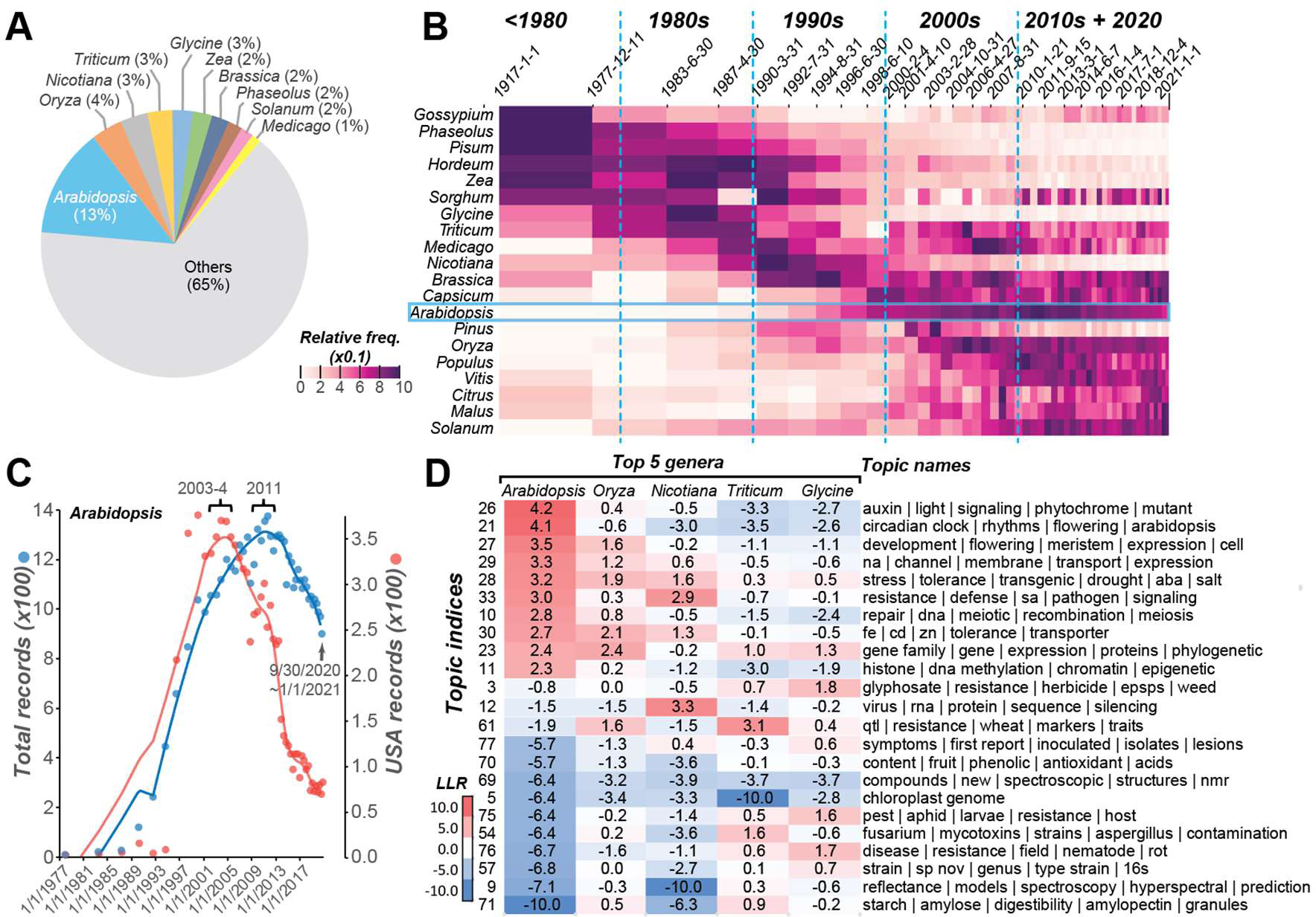
Prevalence of different plant genera in the plant science records. **(A)** Percentage of records mentioning specific genera. **(B)** Change in the prevalence of genera in plant science records over time. **(C)** Changes in the numbers of all records (blue) and records from the US (red) mentioning Arabidopsis over time. The lines are locally weighted scatterplot smoothing fits with fraction parameter=0.2. **(D)** Topical over (red) and under (blue) representation among five genera with the most plant science records. LLR: log 2 likelihood ratios of each topic in each genus.

Assuming the number of publications reflects the intensity of research activities, one hypothesis for the general decline in Arabidopsis research is that advances in, for example, plant transformation, genetic manipulation, and genome research have allowed the adoption of more previously non-model taxa. Consistent with this, there was a precipitous increase in the number of genera being published in the mid-90s to early 2000s during which approaches for plant transgenics became established (*30*), but the number has remained steady since then (**fig. S7A**). The decline in Arabidopsis research is also negatively correlated with the timing of an increase in the number of draft genomes (**fig. S7B, Data S6**). It is plausible that genome availability for other species may have contributed to a shift away from Arabidopsis. By considering both taxa information and research topics, we can identify clear differences in the topical areas preferred by researchers using different plant taxa (**Fig. 4D, Data S7**). For example, studies in auxin/light signaling, the circadian clock, and flowering tend to be carried out in Arabidopsis, while quantitative genetic studies of disease resistance tend to be done in wheat and rice, glyphosate research in soybean, and RNA virus research in tobacco. Taken together, joint analyses of topics and species revealed additional details on preferred models over time, and the preferred topical areas for different taxa.

### Countries differ in their contributions to plant science and topical preference

We next investigated whether there were geographical differences in topical preference among countries by inferring country information from 330,187 records (see **Methods**). The 10 countries with the most records account for 73% of the total, with China and the US contributing to ∼18% each (**Fig. 5A**). The exponential growth in plant science records (green line, **Fig. 1A**) was in large part due to the rapid rise in annual record numbers in China and India (**Fig. 5B**). On the other hand, the US, Japan, Germany, France, and Great Britain had slower rates of growth compared with all non-top 10 countries. The rapid increase in records from China and India was accompanied by a rapid increase in metrics measuring journal impact (**Fig. 5C, fig. S8, Data S6**). For example, using citation score (**Fig. 5C**, see **Methods**), we found that during a 22-year period China (dark green) and India (light green) approached the global average (y=0, yellow) rapidly while some of the other top 10 countries, particularly the US (red) and Japan (yellow green), showed signs of decrease (**Fig. 5C**). It remains to be determined whether these geographical trends reflect changes in priority, investment, and/or interest in plant science research.

**Fig. 5.**
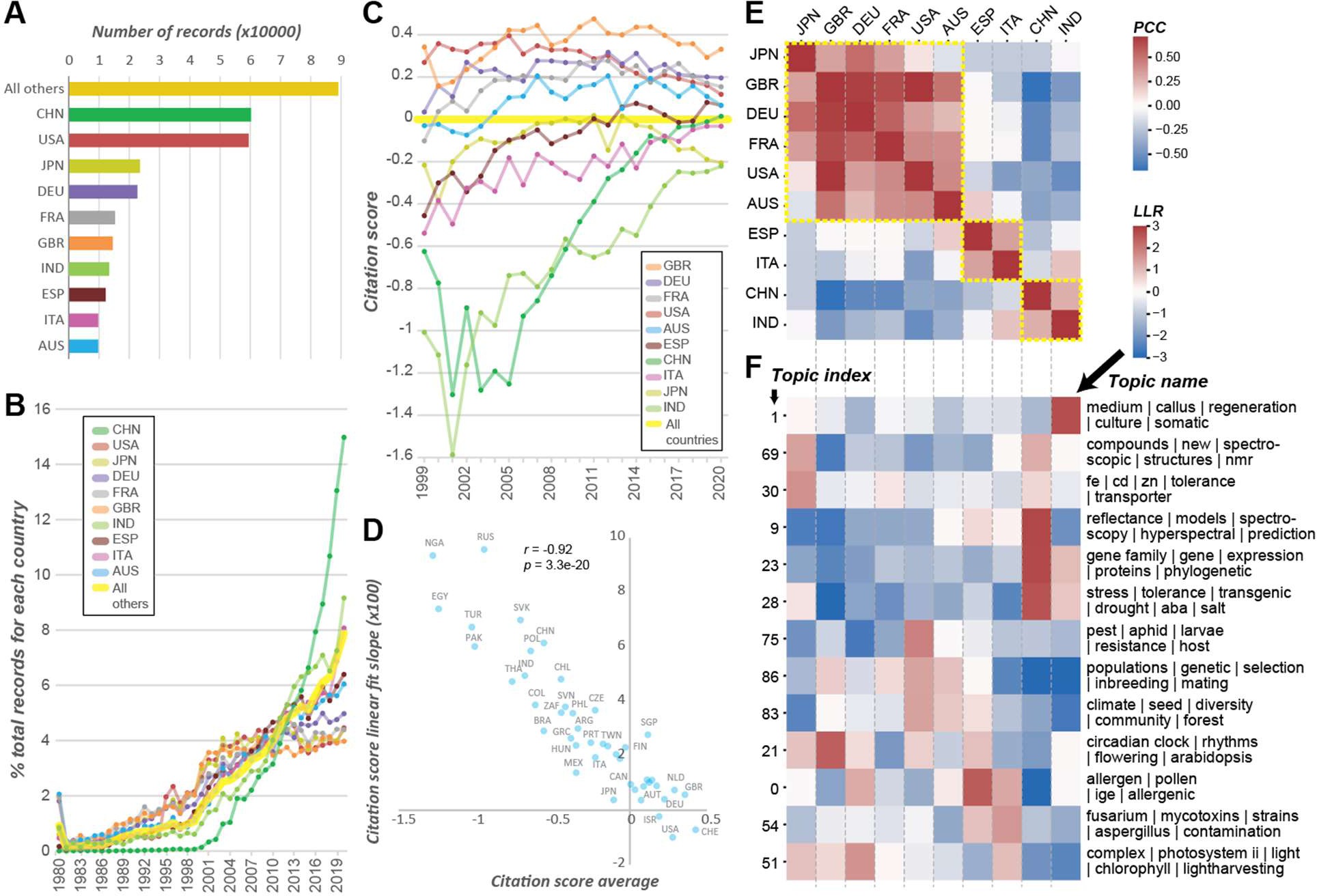
Plant science record numbers, citation scores, and topical preference of different countries. **(A)** Numbers of plant science records for countries with the 10 highest numbers. **(B)** Percentage of all records from each country between 1980 to 2020 for the top 10 countries. **(C)** Difference in citation scores between 1999 to 2020 for the top 10 countries. **(D)** Shown for each country is the relationship between the citation scores averaged between 1999 to 2020 and the slope of linear fit with year as the predictive variable and citation score as the response variable. The countries with >400 records and with <10% missing impact values are included. **(E)** Correlation in topic enrichment scores between the top 10 countries. PCC: Pearson’s Correlation Coefficient. **(F)** Enrichment scores (LLR: Log Likelihood Ratio) of selected topics among the top 10 countries. Red: over-representation, blue: under-representation. Yellow rectangle: countries with more similar topical preferences.

Interestingly, the growth/decline in citation scores over time (measured as the slope of linear fit of year vs. citation score) was significantly and negatively correlated with average citation score (**Fig. 5D**). That is, countries with lower overall metrics tended to experience the strongest increase in citation scores over time. Thus, countries that did not originally have a strong influence on plant sciences now have increased impact. These patterns were also observed when using H-index or journal rank as metrics (**fig. S8, Data S8**) and were not due to increased publication volume, as the metrics were normalized against numbers of records from each country (see **Methods**). In addition, the fact that different metrics with different caveats and assumptions yielded consistent conclusions indicates the robustness of our observations. We also assessed how the plant research foci of countries differ by comparing topical preference (i.e., the degree of enrichment of plant science records in different topics) between countries. For example, Italy and Spain cluster together (yellow rectangle, **Fig. 5E**) partly because of similar research focusing on allergens (topic 0) and mycotoxins (topic 54) and less emphasis on gene family (topic 23) and stress tolerance (topic 28) studies (**Fig. 5F**). There are substantial differences in topical focus between countries (**fig. S9**). For example, research on new plant compounds associated with herbal medicine (topic 69) is a focus in China but not the US, but the opposite is true for population genetics and evolution (topic 86) (**Fig. 5F**). In addition to revealing how plant science research has evolved over time, topic modeling provides additional insights into differences in research foci among different countries, which are informative for science policy considerations.

## DISCUSSION

In this study, topic modeling revealed clear transitions among research topics, which represent paradigm shifts in plant sciences. One limitation of our study is the bias in the PubMed-based corpus. The cellular, molecular, and physiological aspects of plant sciences are well represented, but there are many fewer records related to evolution, ecology, and systematics. Another limitation is the need to assign only one topic to a record when a study is interdisciplinary and straddles multiple topics. Furthermore, a limited number of large, inherently heterogeneous topics were summarized to provide a more concise interpretation that undoubtedly underrepresents the diversity of plant science research. Despite these limitations, dynamic topic modeling revealed changes in plant science research trends that coincide with paradigm shifts in biological science.

Because the key terms defining the topics frequently describe various technologies, this analysis highlights how technological innovation has driven research in the plant sciences, a finding that likely holds for other scientific disciplines. We also found that the pattern of topic evolution is similar to that of succession, where older topics mostly do not persist but appear to be replaced by newer ones. One example is the rise of transcriptome-related topics and the correlated, reduced focus on regulation at levels other than transcription. This raises the question of whether research driven by technology occurs at the detriment of key areas of research where high-throughput studies remain challenging. By analyzing the country of origin, we found that China and India have been the two major contributors to the growth in the plant science records in the last 20 years. Our findings also show an equalizing trend in global plant science where countries without a strong plant science publication presence have had an increased impact over the last 20 years. In addition, we identified significant differences in research topics between countries reflecting potential differences in investment and priorities. Such information is important for discerning differences in research trends across countries and can be considered when making policy decisions about research directions.

## Acknowledgments

We thank Maarten Grootendorst for discussions on topic modeling. We also thank Stacey Harmer, Eva Farre, Ning Jiang, and Robert Last for discussion on their respective research fields and input on how to improve this study and Rudiger Simon for the suggestion to examine differences between countries. We also thank Mae Milton, Christina King, Edmond Anderson, Jingyao Tang, Brianna Brown, Kenia Segura Abá, Eleanor Siler, Thilanka Ranaweera, Huan Chen, Rajneesh Singhal, Paulo Izquierdo, Jyothi Kumar, Daniel Shiu, Elliott Shiu, and Wiggler Catt for their good ideas, personal and professional support, collegiality, fun at parties, as well as the trouble they have caused, which helped us improve as researchers, teachers, mentors, and parents.

## Author contributions

Conceptualization: SHS, MDL

Methodology: SHS

Investigation: SHS

Visualization: SHS, MDL

Funding acquisition: SHS, MDL

Project administration: SHS, MDL

Supervision: SHS

Writing – original draft: SHS

Writing – review & editing: SHS, MDL

## Data and materials availability

The plant science corpus data is available through Zenodo (pending). The codes for topic modeling are available through GitHub (https://github.com/ShiuLab/plant_sci_hist).

## Supplemental Information

**Fig. S1.**
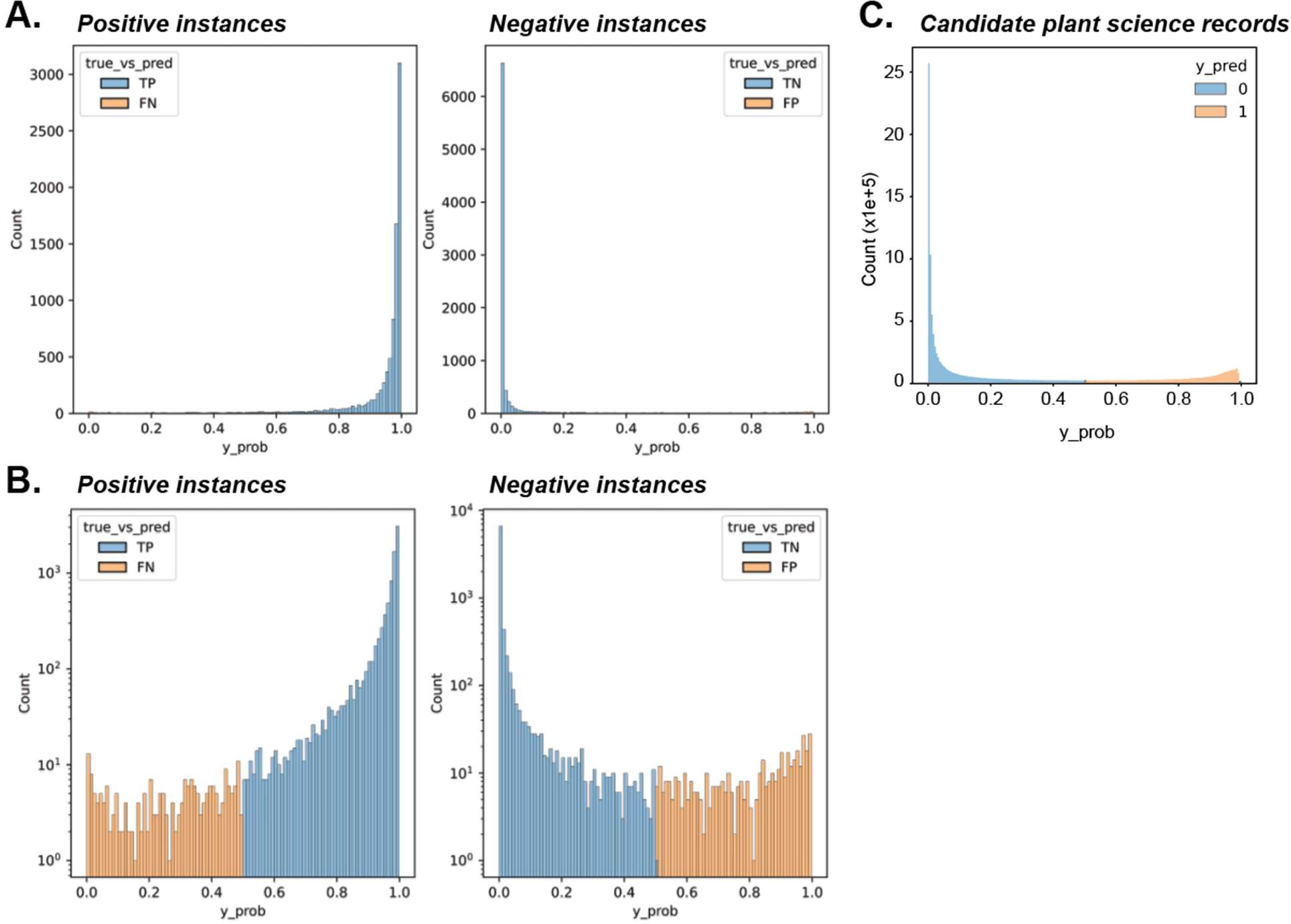
Plant science record classification model performance. **(A)** Prediction probabilities (y_prob) of positive (plant science records) and negative (non-plant science records) instances color coded if they are correctly (blue) or incorrectly (orange) predicted. **(B)** Log-transformed x-axes for the same distributions in **(A)** for better visualization of incorrect predictions. **(C)** Prediction probability distribution for candidate plant science records.

**Fig. S2.**
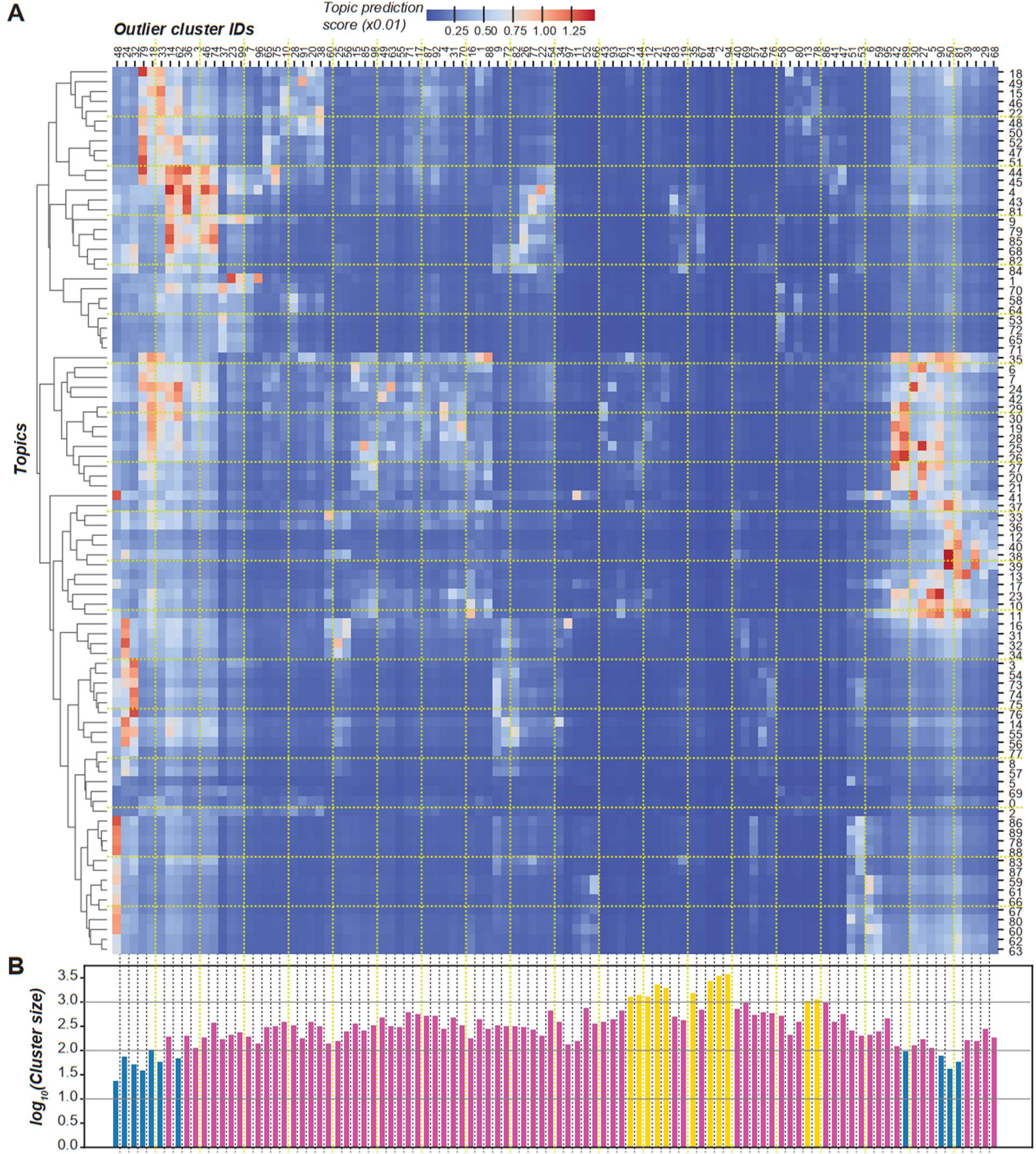
Relationships between outlier clusters and the 90 topics. **(A)** Heatmap demonstrating that some outlier clusters tend to have high prediction scores for multiple topics. Each cell shows the average prediction score of a topic for records in an outlier cluster. **(B)** Size of outlier clusters.

**Fig. S3.**
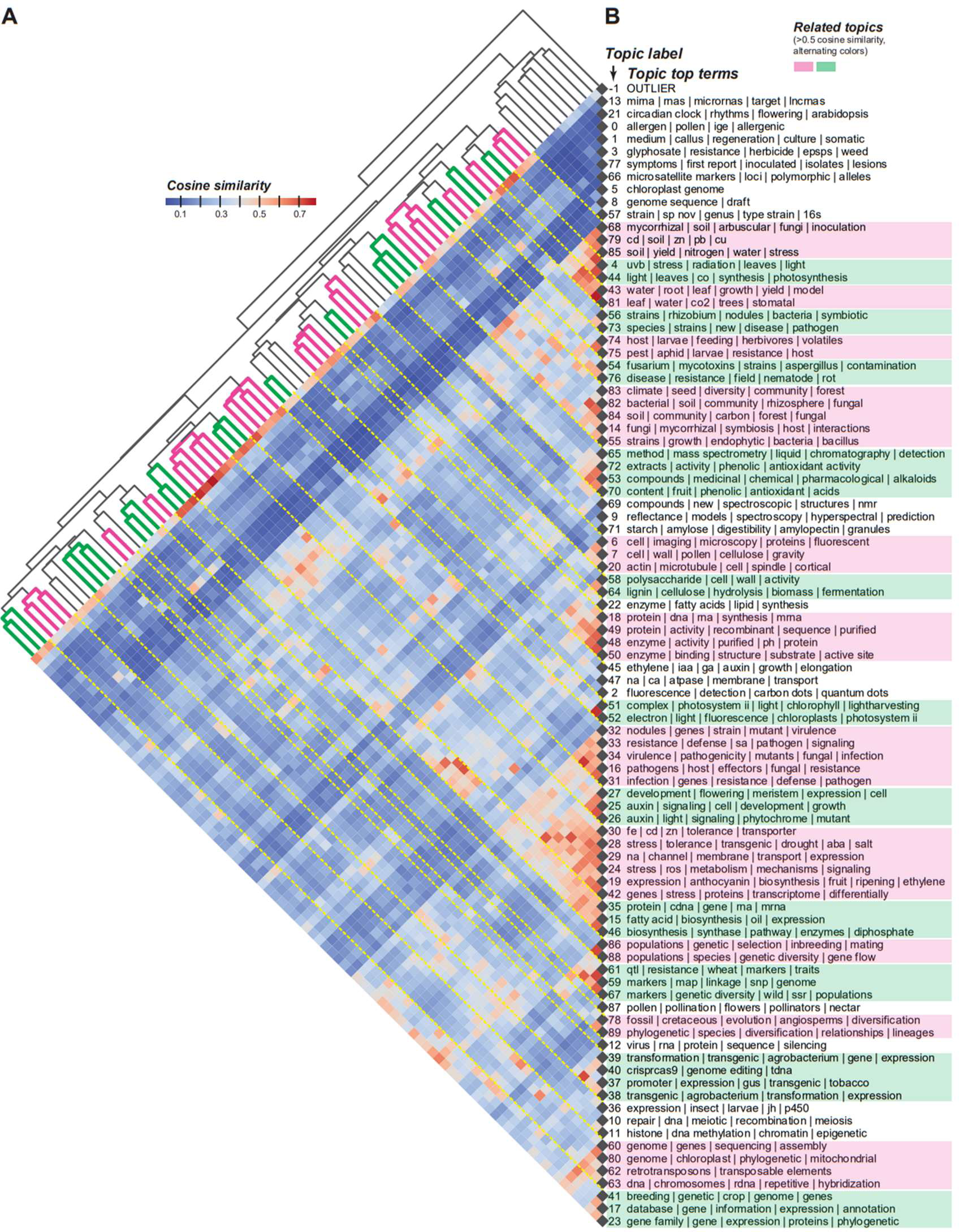
Cosine similarities between topics. **(A)** Heatmap showing cosine similarities between topic pairs. Top-left: hierarchical clustering of the cosine similarity matrix with the Ward algorithm. The branches are colored to indicate groups of related topics. **(B)** Topic labels and names. The topic ordering was based on hierarchical clustering of topics. Colored rectangle: neighboring topics with >0.5 cosine similarities.

**Fig. S4.**
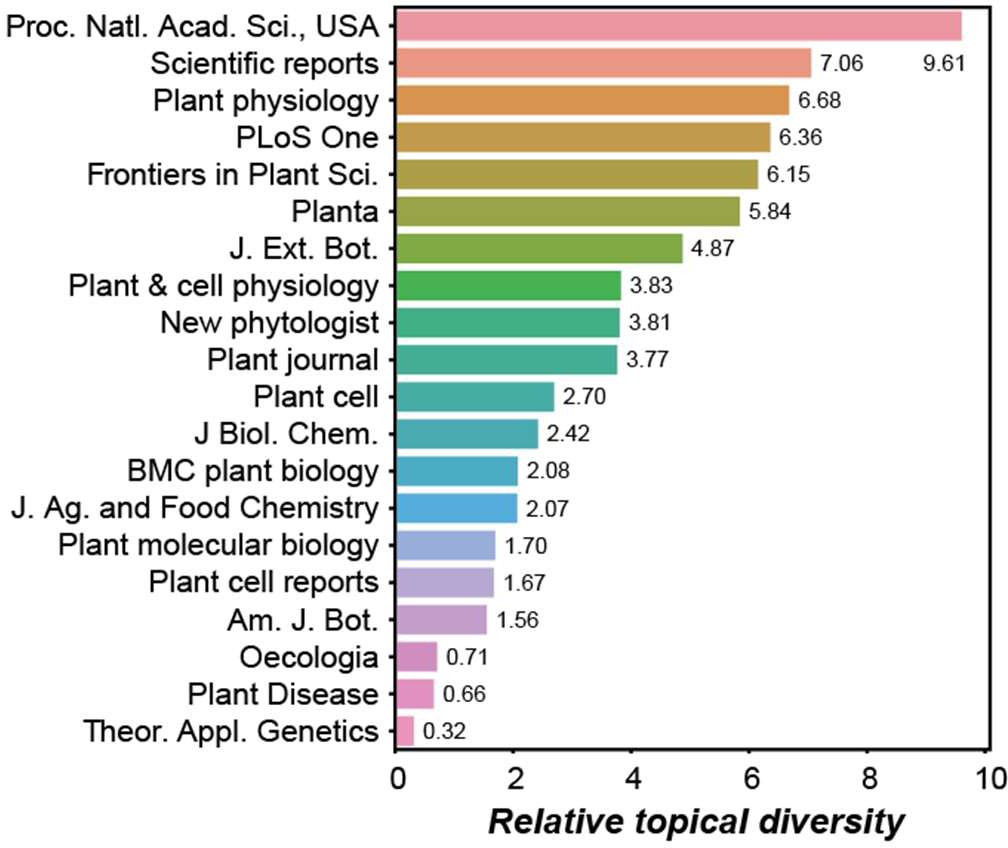
Relative topical diversity for 20 journals. The 20 journals with the most plant science records are shown. The journal names were taken from the journal list in PubMed (https://www.nlm.nih.gov/bsd/serfile_addedinfo.html).

**Fig. S5.**
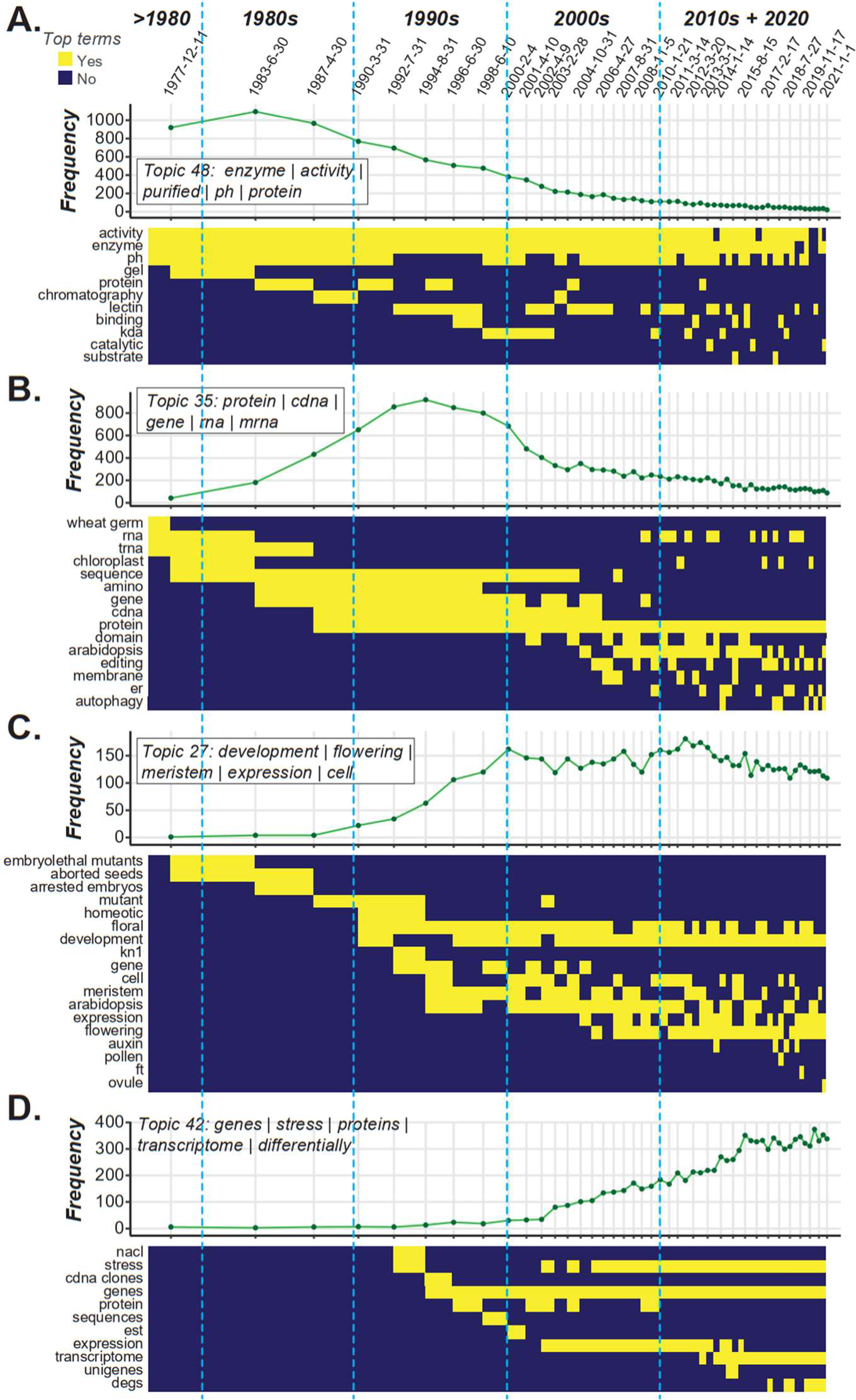
Topical frequency and top terms during different time periods. **(A-D)** Different patterns of topical frequency distributions for example topics **(A)** 48, **(B)** 35, **(C)** 27, and **(D)** 42. For each topic, the top graph shows the frequency of topical records in each time bin, which are the same as those in Fig. 3 (green line), and the end date for each bin is indicated. The heatmap below each line plot depicts whether a term is among the top terms in a time bin (yellow) or not (blue). Blue dotted lines delineate different decades.

**Fig. S6.**
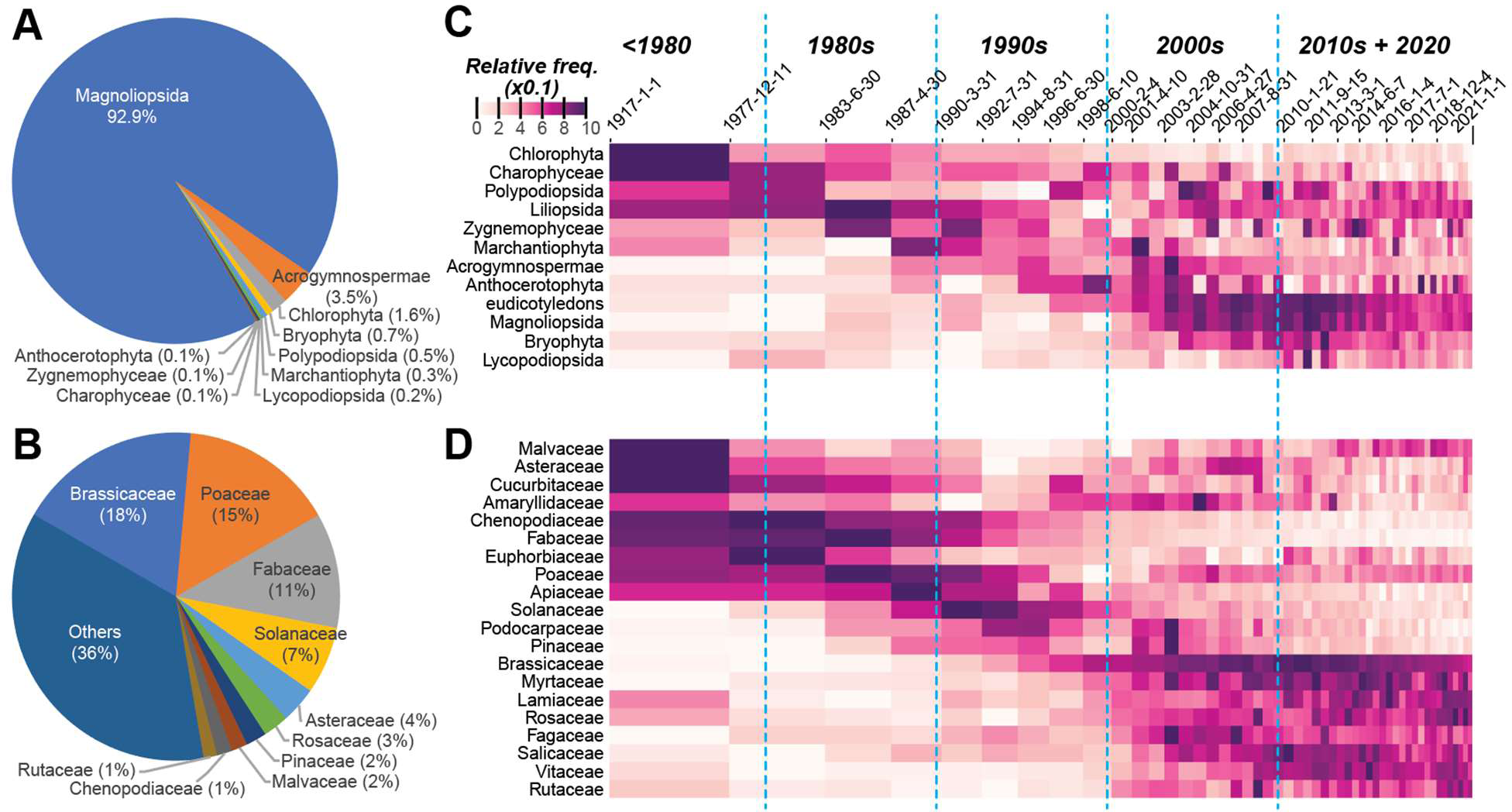
Prevalence of records mentioning different taxonomic groups in Viridiplantae. **(A,B)** Percentage of records mentioning specific taxa at the (**A)** major lineage and **(B)** family levels. **(C,D)** The prevalence of taxon mentions over time at the **(C)** major lineage and **(E)** family levels.

**Fig. S7.**
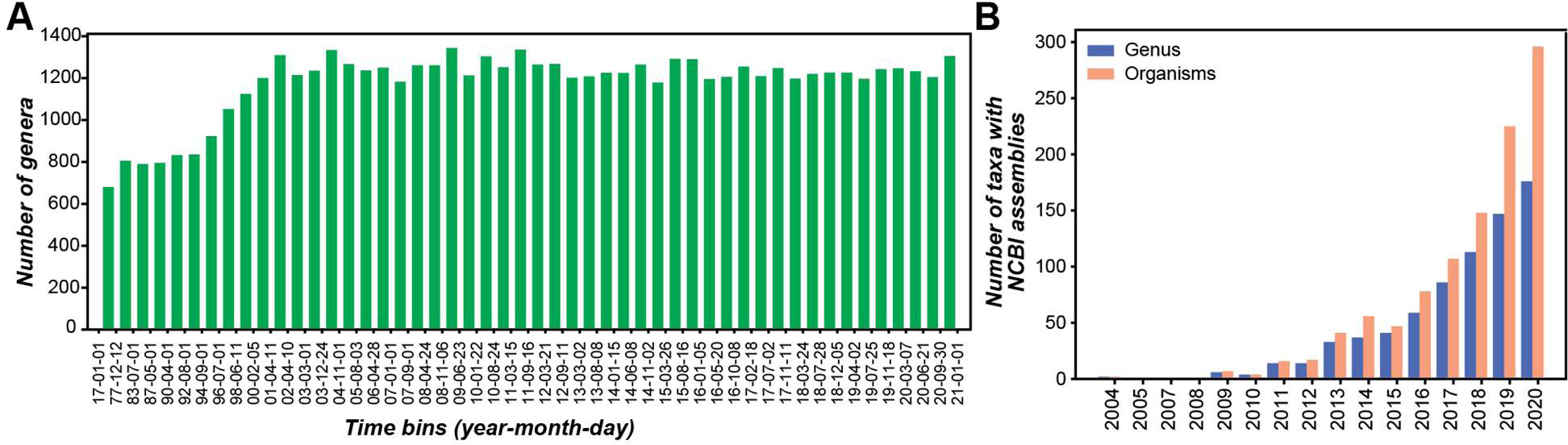
Changes in the numbers of taxa reported over time. **(A)** Number of genera being mentioned in plant science records during different time bins (the date indicates the end date of that bin, exclusive). **(B)** Numbers of genera (blue) and organisms (salmon) with draft genomes available from National Center of Biotechnology Information in different years.

**Fig. S8.**
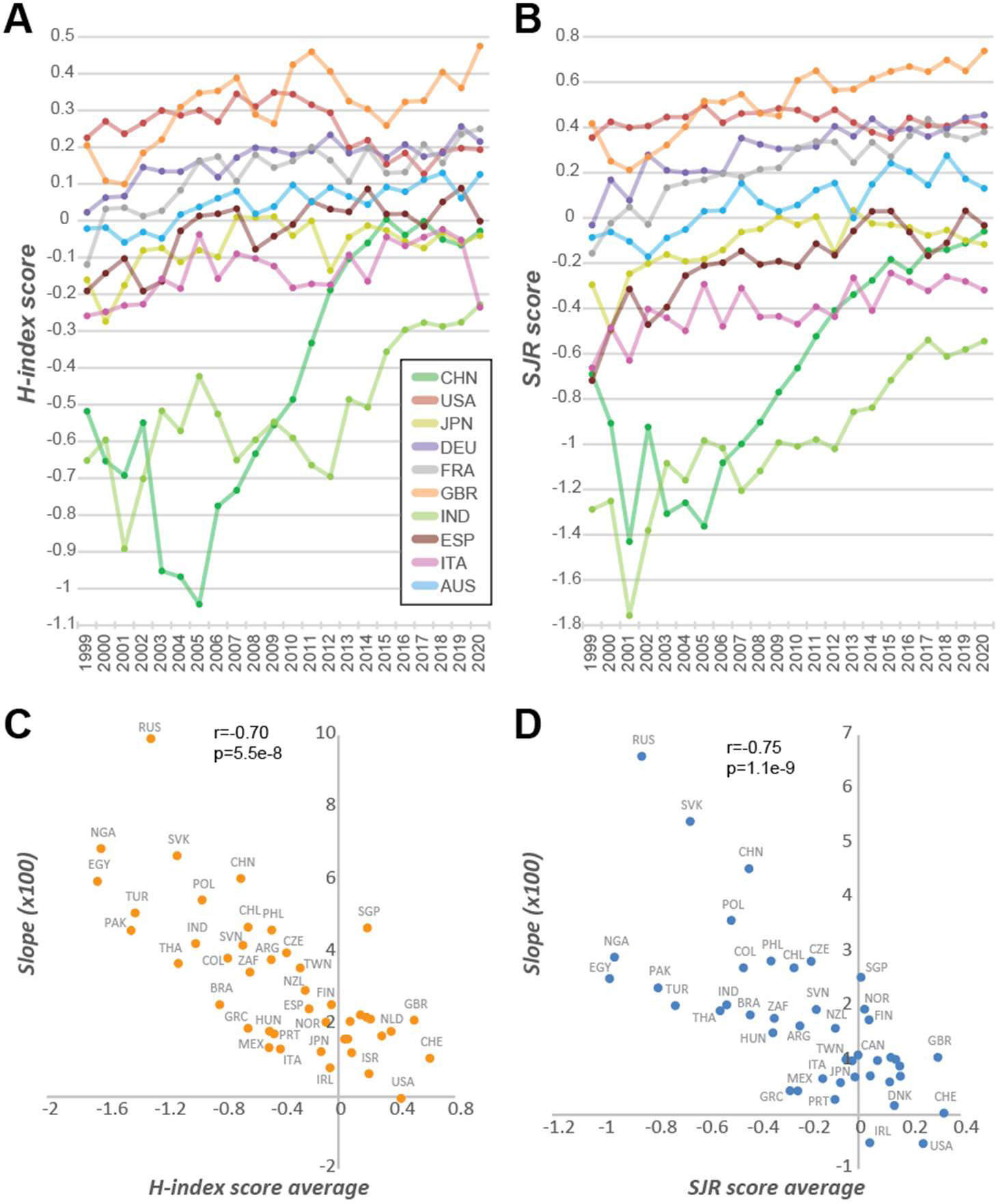
Change in country impact on plant science over time. **(A,B)** Difference in two impact metrics from 1999 to 2020 for the 10 countries with the highest number of plant science records. **(A)** H-index. **(B)** SCImago Journal Rank (SJR). **(C,D)** Plots show the relationships between the impact metrics (H-index in **(C)**, SJR in **(D)**) averaged between 1999 to 2020 and the slopes of linear fits with years as the predictive variable and impact metric as the response variable for different countries (A3 country codes shown). The countries with >400 records and with <10% missing impact values are included.

**Fig. S9.**
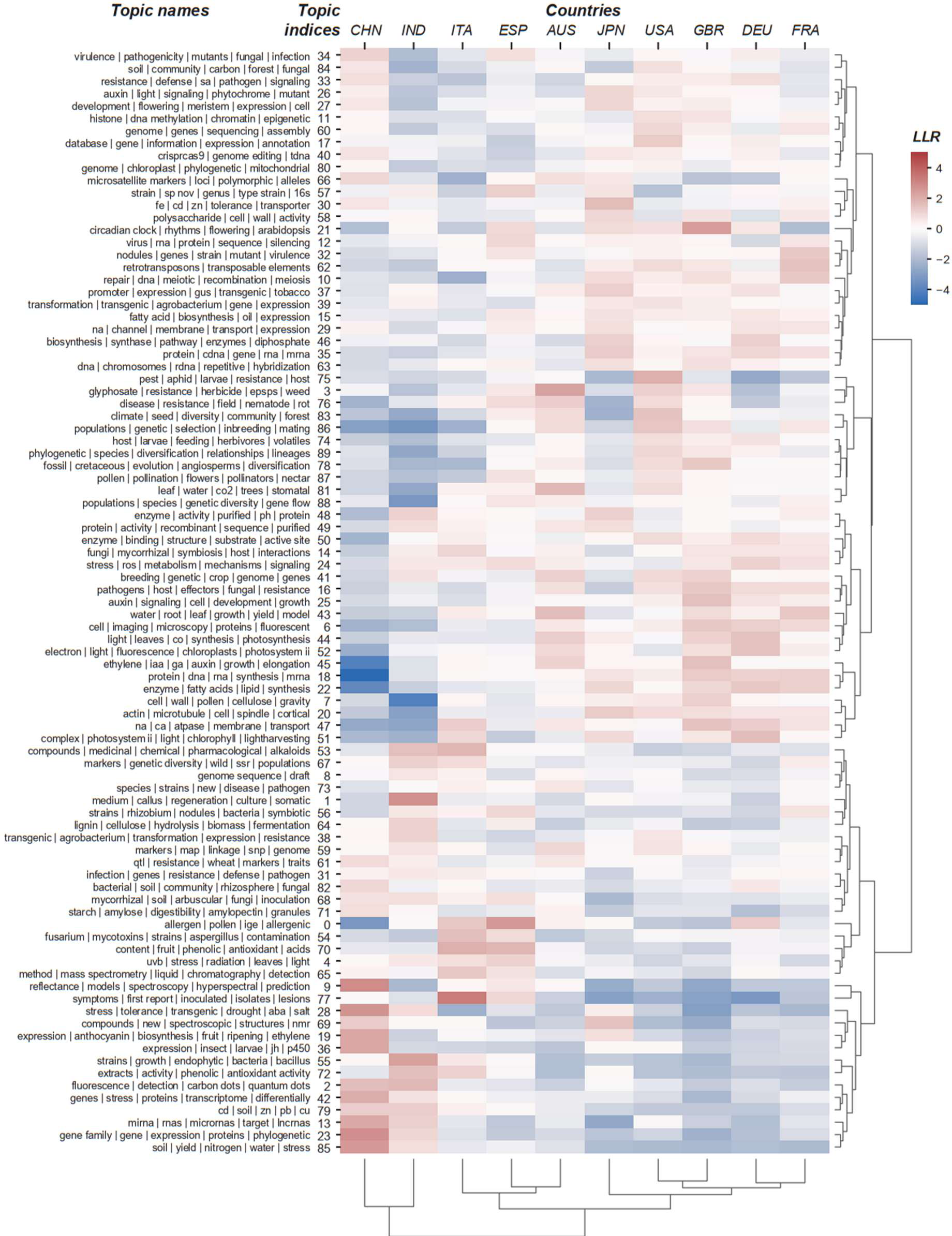
Country topical preference. Enrichment scores (LLR: Log Likelihood Ratio) of topics among the top 10 countries. Red: over-representation, blue: under-representation.

### Data S1. Summary of source journals for plant science records, prediction models, and top Tf-Idf features

- Sheet – Candidate plant sci record j counts: Number of records from each journal in the candidate plant science corpus (before classification)
- Sheet - Plant sci record j count: Number of records from each journal in the plant science corpus (after classification)
- Sheet – Model summary: Model type, text used (txt_flag), and model parameters used
- Sheet - Model performance: Performance of different model and parameter combinations on the validation data set
- Sheet – Tf-Idf features: The average SHAP values of Tf-Idf (Term frequency-Inverse document frequency) features associated with different terms

### Data S2. Numbers of records in topics identified from preliminary topic models

- Sheet – Topics generated with a model based on BioBERT embeddings
- Sheet – Topics generated with a model based on distilBERT embeddings
- Sheet – Topics generated with a model based on SciBERT embeddings

### Data S3. Final topic model labels and top terms for topics

- Sheet – Topic label: The topic index and top 10 terms with the highest cTf-Idf values
- Sheets – 0 to 89: The top 50 terms and their c-Tf-Idf values for topics 0 to 89.

### Data S4. UMAP representations of different topics

For a topic *T*, records in the UMAP graph are colored red and records not in *T* are colored gray.

### Data S5. Timestamps and dates for dynamic topic modeling

Columns are: (1) order index, (2) bin_idx – relative positions of bin labels, (3) bin_timestamp – UNIX time in seconds, and (4) bin_date – month/day/year.

### Data S6. Change in taxon record numbers and genome assemblies available over time

- Sheet – Genus: Number of records mentioning a genus during different time periods (in Unix timestamp) for the top 100 genera.
- Sheet – Genus: Number of records mentioning a family during different time periods (in Unix timestamp) for the top 100 families.
- Sheet – Genus: Number of records mentioning an order during different time periods (in Unix timestamp) for the top 20 orders.
- Sheet – Special levels: Number of records mentioning 12 selected taxonomic levels higher than the order level during different time periods (in Unix timestamp).
- Sheets – Genome assembly: Plant genome assemblies available from NCBI as of 10/28/22.

### Data S7. Taxon topical preference

- Sheet – 5 genera LLR: The log likelihood ratio of each topic in each of the top five genera with the highest numbers of plant science records
- Sheets – 5 genera: For each genus, the columns are: (1) topic, (2) the Fisher’s Exact Test p-value (Pvalue), (3-6) numbers of records in topic *T* and in genus *X* (n_inT_inX), in *T* but not in *X* (n_inT_niX), not in *T* but in *X* (n_niT_inX), and not in *T* and *X* (n_niT_niX) that were used to construct 2x2 tables for the tests, and (7) the log likelihood ratio generated with the 2x2 tables.

### Data S8. Impact metrics of countries in different years

- Sheet – prank: Journal percentile rank scores for countries (three letter country codes following https://www.iban.com/country-codes) in different years from 1999 to 2020.
- Sheet – sjr: Scimago Journal rank scores
- Sheet – hidx: H-Index scores
- Sheet – cite: Citation scores

## Supplemental Methods

### Collection and pre-processing of a candidate plant science corpus

For reproducibility purposes, a random state value of 20220609 (project start date) was used throughout the study. The PubMed baseline files containing citation information (ftp://ftp.ncbi.nlm.nih.gov/pubmed/baseline/) were downloaded on 11/11/2021. To narrow down the records to plant science-related citations, a candidate citation was identified as having at least one of the keywords “plant,” “plants,” “botany,” “botanical,” “planta,” and “plantarum” (and their corresponding upper case and plural forms), or plant taxon identifiers from NCBI Taxonomy (https://www.ncbi.nlm.nih.gov/taxonomy) or USDA PLANTS Database (https://plants.sc.egov.usda.gov/home). The taxon identifiers include all taxon names including and at taxonomic levels below “Viridiplantae” till the genus level (species names not used). This led to 51,395 keywords. After the keyword search, qualified entries were removed if they were duplicated, lacked titles and/or abstracts, or were corrections, errata, or withdrawn articles. This left 1,385,417 citations, which were considered the candidate plant science corpus (i.e., a collection of texts). For further analysis, the title and abstract for each citation were combined into a single entry. Text was pre-processed by lowercasing, removing stop-words (i.e., common words), removing non-alphanumeric and non-white space characters (except Greek letters, dashes, and commas), and applying lemmatization (i.e., grouping inflected forms of a word as a single word) for comparison. Because lemmatization led to truncated scientific terms, it was not included in the final pre-processing pipeline.

### Definition of positive/negative examples

Upon closer examination, a large number of false positives were identified in the candidate plant science records. To further narrow down citations with a plant science focus, text classification was used to distinguish plant science and non-plant science articles (see next section). For the classification task, a negative set (i.e., non-plant science citations) was defined as entries from 7,360 journals that appeared <20 times in the filtered data (total=43,329, journal candidate count, **Data S1**). For the positive examples (i.e., true plant science citations), 43,329 plant science citations (positive examples) were sampled from 17 established plant science journals each with >2,000 entries in the filtered dataset: ‘Plant physiology’, ‘Frontiers in plant science’, ‘Planta’, ‘The Plant journal : for cell and molecular biology’, ‘Journal of experimental botany’, ‘Plant molecular biology’, ‘The New phytologist’, ‘The Plant cell’, ‘Phytochemistry’, ‘Plant & cell physiology’, ‘American journal of botany’, ‘Annals of botany’, ‘BMC plant biology’, ‘Tree physiology’, ‘Molecular plant-microbe interactions : MPMI’, ‘Plant biology’, and ‘Plant biotechnology journal’ (journal candidate count, **Data S1**). Plant biotechnology journal was included, but only 1894 records remained after removal of duplicates, articles with missing info, and/or withdrawn articles. The positive and negative sets were randomly split into training and testing subsets (4:1) while maintaining a 1:1 positive-to-negative ratio.

### Text classification based on Tf and Tf-Idf

Instead of using the pre-processed text as features for building classification models directly, text embeddings (i.e., representations of texts in vectors) were used as features. These embeddings were generated using four approaches (model summary, **Data S1**): Term-frequency (Tf), Tf-Inverse document frequency (Tf-Idf, (*31*)), Word2Vec (*32*), and Bidirectional Encoder Representations from Transformers (BERT, (*6*)). The Tf- and Tf-Idf-based features were generated with CountVectorizer and TfidfVectorizer, respectively, from Scikit-Learn (*33*). Different maximum features (1e4 to 1e5) and n-gram ranges (uni-, bi-, and tri-grams) were tested. The features were selected based on the *p-*value of Chi-square tests testing whether a feature had a higher-than-expected value among the positive or negative classes. Four different *p-*value thresholds were tested for feature selection. The selected features were then used to retrain vectorizers with the pre-processed training texts to generate feature values for classification. The classification model used was XGBoost (*34*) with five combinations of the following hyperparameters tested during five-fold stratified cross-validation: min_child_weight=[1, 5, 10], gamma=[0.5, 1, 1.5, 2,5], subsample=[0.6, 0.8, 1.0], colsample_bytree=[0.6, 0.8, 1.0], and max_depth=[3, 4, 5]. The rest of the hyperparameters were held constant: learning_rate=0.2, n_estimators=600, objective=binary:logistic. RandomizedSearchCV from Scikit-Learn was used for hyperparameter tuning and cross-validation with scoring=F1-score.

Because the Tf-Idf model had a relatively high model performance and was relatively easy to interpret (terms are frequency-based, instead of embedding-based like those generated by Word2Vec and BERT), the Tf-Idf model was selected as input to SHapley Additive exPlanations (SHAP, (*35*)) to assess the importance of terms. Because the Tf-Idf model was based on XGBoost, a tree-based algorithm, the TreeExplainer module in SHAP was used to determine a SHAP value for each entry in the training dataset for each Tf-Idf feature. The SHAP value indicates the degree to which a feature positively or negatively affects the underlying prediction. The importance of a Tf-Idf feature was calculated as the average SHAP value of that feature among all instances. Because a Tf-Idf feature is generated based on a specific term. The importance of the Tf-Idf feature indicates the importance of the associated term.

### Text classification based on Word2Vec

The preprocessed texts were first split into train, validation, and test subsets (8:1:1). The texts in each subset were converted to three n-gram lists: a unigram list obtained by splitting tokens based on the space character, or bi- and tri-gram lists built with Gensim (*36*). Each n-gram list of the training subset was next used to fit a Skip-gram Word2Vec model with vector_size=300, window=8, min_count=(5, 10, or 20), sg=1, and epochs=30. The Word2Vec model was used to generate word embeddings for train, validate, and test subsets. In the meantime, a tokenizer was trained with train subset unigrams using Tensorflow (*37*) and used to tokenize texts in each subset and turn each token into indices to use as features for training text classification models. To ensure all citations had the same number of features (500), longer texts were truncated, and shorter ones were zero-padded. A deep learning model was used to train a text classifier with an input layer the same size as the feature number, an attention layer incorporating embedding information for each feature, two bi-directional Long-Short-Term-Memory layers (15 units each), a dense layer (64 units), and a final, output layer with 2 units. During training, adam, accuracy, and sparse_categorical_crossentropy were used as the optimizer, evaluation metric, and loss function, respectively. The training process lasted 30 epochs with early stopping if validation loss did not improve in five epochs. An F1 score was calculated for each n-gram list and min_count parameter combination to select the best model (model summary, **Data S1**).

### Text classification based on BERT models

Two pre-trained models were used for BERT-based classification: DistilBERT (Huggingface repository (*38*) model name and version: distilbert-base-uncased, (*39*)) and SciBERT (allenai/scibert-scivocab-uncased, (*13*). In both cases, tokenizers were re-trained with the training data. BERT-based models had the following architecture: the token indices (512 values for each token) and associated masked values as input layers, pretrained BERT layer (512x768) excluding outputs, a 1D pooling layer (768 units), a dense layer (64 units), and an output layer (2 units). The rest of the training parameters were the same as those for Word2Vec-based models, except training lasted for 20 epochs. Cross-validation F1-scores for all models were compared and used to select the best model for each feature extraction method, hyperparameter combination, and modeling algorithm or architecture (model summary, **Data S1**). The best model was the Word2Vec-based model (min_count=20, window=8, ngram=3), which was applied to the candidate plant science corpus to identify a set of plant science citations for further analysis. The candidate plant science records predicted as being in the positive class (421,658) by the model were collectively referred to as the “plant science corpus.”

### Plant science record classification

In PubMed, 1,384,718 citations containing “plant” or any plant taxon names (from the phylum to genus level) were considered candidate plant science citations. To further distinguish plant science citations from those in other fields, text classification models were trained using titles and abstracts of positive examples consisting of citations from 17 plant science journals, each with >2,000 entries in PubMed, and negative examples consisting of records from journals with fewer than 20 entries in the candidate set. Among four models tested (see **Methods**), the best model (built with Word2Vec embeddings) had a cross validation F1 of 0.964 (random guess F1=0.5, perfect model F1=1, **Data S1**). When testing the model using 17,330 testing set citations independent from the training set, the F1 remained high at 0.961 (false negative rate=2.6%, false positive rate=5.2%, **fig S1A,B**).

To better understand how the models classified plant science articles, we identified important terms from a more easily interpretable model (Term frequency-Inverse document frequency, Tf-Idf, model; F1=0.934) using Shapley Additive Explanations (*35*); 136 terms contributed to predicting plant science records (e.g., Arabidopsis, xylem, seedling) and 138 terms contributed to non-plant science record predictions (e.g., patients, clinical, mice; Tf-Idf feature sheet, **table S1**). Applying the best model to the candidate plant science records led to 421,658 positive predictions, hereafter referred to as “plant science records” (**fig. S1C**, **Data S1**).

### Global topic modeling

BERTopic (*40*) was used for preliminary topic modeling with n-grams=(1,2) and with an embedding initially generated by DistilBERT, SciBERT, or BioBERT (dmis-lab/biobert-base-cased-v1.2, (*41*)). The embedding models converted pre-processed texts to embeddings. The topics generated based on the three embeddings were similar (**Data S2**). However, SciBERT-, BioBERT-, and distilBERT-based embedding models had different numbers of outlier records (268,848, 293,790, and 323,876, respectively) with topic index=-1. In addition to generating the fewest outliers, the SciBERT-based model led to the highest number of topics. Therefore, SciBERT was chosen as the embedding model for the final round of topic modeling. Modeling consisted of three steps. First, document embeddings were generated with SentenceTransformer (*42*). Second, a clustering model to aggregate documents into clusters using hdbscan (*43*) was initialized with min_cluster_size=500, metric=euclidean, cluster_selection_method=eom, min_samples=5. Third, the embedding and the initialized hdbscan model were used in BERTopic to model topics with neighbors=10, nr_topics=500, ngram_range=(1,2). Using these parameters, 90 topics were identified. The initial topic assignments were conservative, and 241,567 records were considered outliers (i.e., documents not assigned to any of the 90 topics). After assessing the prediction scores of all records generated from the fitted topic models, the 95-percentile score was 0.0155. This score was used as the threshold for assigning outliers to topics: if the maximum prediction score was above the threshold and this maximum score was for topic *t*, then the outlier was assigned to *t*. After the re-assignment, 49,228 records remained outliers. To assess if some of the outliers were not assigned because they could be assigned to multiple topics, the prediction scores of the records were used to put records into 100 clusters using *k-*means. Each cluster was then assessed to determine if the outlier records in a cluster tended to have higher prediction scores across multiple topics (**fig. S2**).

### Topics that are most and least well-connected to other topics

The most well-connected topics in the network include topic 24 (stress mechanisms, median cosine similarity=0.36), topic 42 (genes, stress, and transcriptomes, 0.34), and topic 35 (molecular genetics, 0.32, all t-test *p*-values<1e-22). The least connected topics include topic 0 (allergen research, median cosine similarity=0.12), topic 21 (clock biology, 0.12), topic 1 (tissue culture, 0.15), and topic 69 (identification of compounds with spectroscopic methods, 0.15; all t-test *p-*values<1e-24). Topics 0, 1, and 69 are specialized topics, it is surprising that topic 21 is not as well connected as explained in the main text.

### Analysis of documents based on the topic model

From the fitted topic model, we obtained a topic-by-term matrix with 91 rows, where each row corresponded to a topic or outlier, and 1.85e7 columns, where each column corresponded to a term in the plant science record vocabulary. This topic-by-term matrix was filled with c-Tf-Idf values for each topic-term combination. The similarity between each pair of topics, excluding the outlier rows, was determined by determining the cosine similarity using the c-Tf-Idf vectors of the two topics in question (**fig. S4**). Topics with cosine similarities ≥0.6 were connected in the network graph shown in **Fig. 2**. The relative topical diversity of a journal *j* (*v*_j_) was calculated as follows:

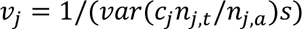

Where *var* is variance, *c*_j_ is the median cosine similarity between a topic *t* and non-*t* topics, *n*_j,t_ is the number of records of journal *j* in topic *t*, *n*_j,a_ is the number of all records of journal *j* in the plant science corpus, and *s* is a scaling factor, which is set to 1e4 so that the relative topical diversity value is between 0 and 10 for a more straightforward comparison. The top 50 terms for each topic are available in **Data S3**. To examine the relationships between documents in a scatter plot with two dimensions, we needed to reduce the dimensionality of BERT embeddings (containing information on how two documents were related to each other) from 768 to two. To accomplish this, Uniform Manifold Approximation and Projection (UMAP, (*14*)) was used. The UMAP parameters tested were n_neighbors=(5, 10, 15, 40), min_dist=(0, 0.1, 0.25), and metric=(cosine, euclidean, canberra, mahalanobis, correlation). After comparing the graphs generated with different UMAP parameter combinations, we used the parameter combination n_neighbors=24, min_dist=0.1, and metric=cosine to generate the final 2D representation (**Data S4**).

### Topical diversity among top journals with the most plant science records

Using a relative topic diversity measure (ranging from 0 to 10, see **Methods**), we found that there was a wide range of topical diversity among 20 journals with the largest numbers of plant science records (**fig. S3**). The four journals with the highest relative topical diversities are *Proceedings of the National Academy of Sciences, USA* (9.6), *Scientific Reports* (7.1), *Plant Physiology* (6.7), and *PLoS ONE* (6.4). The high diversities are consistent with the broad, editorial scopes of these journals. The four journals with the lowest diversities are *American Journal of Botany* (1.6), *Oecologia* (0.7), *Plant Disease* (0.7), and *Theoretical and Applied Genetics* (0.3), which reflects their discipline-specific focus and audience of classical botanists, ecologists, plant pathologists, and specific groups of geneticists.

### Dynamic topic modeling

The codes for dynamic modeling were based on _topic_over_time.py in BERTopics and modified to allow additional outputs for debugging and graphing purposes. The plant science citations were binned into 50 subsets chronologically. Because the numbers of documents increased exponentially over time, instead of dividing them based on equal-sized time intervals, which would result in fewer records at earlier timepoints and introduce bias, we divided them into time bins of similar size (∼8,400 documents). Thus, the earlier time subsets had larger time spans compared with later time subsets. If equal-size time intervals were used, the numbers of documents between the intervals would differ greatly; the earlier time points would have many fewer records, which may introduce bias. Prior to binning the subsets, the publication dates were converted to UNIX time (timestamp) in seconds; the plant science records start in 1917-11-1 (timestamp=-1646247600.0) and end in 2021-1-1 (timestamp=1609477201). The starting dates and corresponding timestamps for the 50 subsets including the end date are in **Data S5**. The input data included the pre-processed texts, topic assignments of records from global topic modeling, and the binned timestamps of records. Three additional parameters were set for topics_over_time, namely: nr_bin=50 (number of bins), evolution_tuning=True, and global_tuning=False. The evolution_tuning parameter specified that averaged c-Tf-Idf values for a topic be calculated in neighboring time bins to reduce fluctuation in c-Tf-Idf values. The global_tuning parameter was set to False because of the possibility that some non-existing terms could have a high c-Tf-Idf for a time bin simply because there was a high global c-Tf-Idf value for that term.

### Taxa information

To identify a taxon or taxa in all plant science records, NCBI Taxonomy taxdump datasets were downloaded from the NCBI FTP site (https://ftp.ncbi.nlm.nih.gov/pub/taxonomy/new_taxdump/) on 9/20/2022. The highest level taxon was Viridiplantae, and all its child taxa were parsed and used as queries in searches against the plant science corpus. In addition, a species-over-time analysis was conducted using the same time bins as used for dynamic topic models. The number of records in different time bins for top taxa are in the genus, family, order, and additional species level sheet in **Data S6**. The degree of over-/under-representation of a taxon X in a research topic T was assessed using the p-value of a Fisher’s exact test for a 2×2 table consisting of the numbers of records in both X and T, in X but not T, in T but not X, and in neither (**Data S7**).

For analysis of plant taxa with genome information, genome data of taxa in Viridiplantae were obtained from the NCBI Genome data-hub (https://www.ncbi.nlm.nih.gov/data-hub/genome) on 10/28/22. There were 2,384 plant genome assemblies belonging to 1,231 species in 559 genera (genome assembly sheet, **Data S6**). The date of the assembly was used as a proxy for the time when a genome was sequenced. However, some species have updated assemblies and have more recent data than when the genome first became available.

### Taxa being studied in the plant science records

Flowering plants (Magnoliopsida) are found in 93% of records, while most other lineages are discussed in <1% of records, with conifers and related species being exceptions (Acrogynomsopermae, 3.5%, **fig. S6A**). At the family level, the mustard (Brassicaceae), grass (Poaceae), pea (Fabaceae), and nightshade (Solanaceae) families are in 51% of records (**fig. S6B**). The prominence of the mustard family in plant science research is due to the *Brassica* and *Arabidopsis* genera (**Fig. 4A**). When examining the prevalence of taxa being studied over time, clear patterns of turnovers emerged (**Fig. 4B**, **fig. S6C,D**). While the study of monocot species (Liliopsida) has remained steady, there was a significant up-tick in the prevalence of eudicot (eudicotyledon) records in the late 90s (**fig. S6C**), which can be attributed to the increased number of studies in the mustard, myrtle (Myrtaceae), and mint (Lamiaceae) families among others (**fig. S6D**). At the genus level, records mentioning *Gossypium* (cotton), *Phaseolus* (bean), *Hordeum* (wheat), and *Zea* (corn), similar to the topics in the early category, were prevalent till the 80s or 90s but have mostly declined in number since (**Fig. 4B**). In contrast, *Capsicum, Arabidopsis, Oryza, Vitus,* and *Solanum* research has become more prevalent over the last 20 years.

### Geographical information for the plant science corpus

The geographical information (country) of authors in the plant science corpus was obtained from the address (AD) fields of first authors in Medline XML records accessible through the NCBI EUtility API (https://www.ncbi.nlm.nih.gov/books/NBK25501/). Because only first author affiliations are available for records published before December 2014, only the first author’s location was considered to ensure consistency between records before and after that date. Among the 421,658 records in the plant science corpus, 421,585 had Medline records and 421,276 had unique PMIDs. Among the records with unique PMIDs, 401,807 contained address fields. For each of the remaining records, the AD field content was split into tokens with a “,” delimiter, and the token likely containing geographical info (referred to as location tokens) was selected as either the last token or the second to last token if the last token contained “@” indicating the presence of an email address. Because of the inconsistency in how geographical information was described in the location tokens (e.g., country, state, city, zip code, name of institution, and different combinations of the above), the following four approaches were used to convert location tokens into countries.

The first approach was a brute force search where full names and alpha-3 codes of current countries (ISO 3166-1), current country subregions (ISO 3166-2), and historical country (i.e., country that no longer exists, ISO 3166-3) were used to search the address fields. To reduce false positives using alpha-3 codes, a space prior to each code is required for the match. The first approach allowed the identification of 361,242, 16,573, and 279,839 records with current country, historical country, and subregion information, respectively. The second method was the use of a heuristic based on common address field structures to identify “location strings” toward the end of address fields that likely represent countries, then the use of the Python pycountry module to confirm the presence of country information. This approach led to 329,025 records with country information. The third approach was to parse first author email addresses (90,799 records), recover top-level domain information, and use country code Top Level Domain (ccTLD) data from the ISO 3166 Wikipedia page to define countries (72,640 records). Only a subset of email addresses contain country information because some are from companies (.com), non-profit organizations (.org), and others. Because a large number of records with address fields still did not have country information after taking the above three approaches, another approach was implemented to query address fields against a locally installed Nominatim server (v.4.2.3, https://github.com/mediagis/nominatim-docker) using OpenStreetMap data from GEOFABRIK (https://www.geofabrik.de/) to find locations. Initial testing indicated that the use of full address strings led to false positives and the computing resource requirement for running the server was high. Thus, only location strings from the second approach that did not lead to country information were used as queries. Because multiple potential matches were returned for each query, the results were sorted based on their location importance values. The above steps led to an additional 72,401 records with country information.

Examining the overlap in country information between approaches revealed that brute force current country and pycountry searches were consistent 97.1% of the time. In addition, both approaches had high consistency with the email-based approach (92.4% and 93.9%). However, brute force subregion and Nominatim-based predictions had the lowest consistencies with the above three approaches (39.8%–47.9%) and each other. Thus, a record’s country information was finalized if the information was consistent between any two approaches, except between the brute force subregion and Nominatim searches. This led to 330,328 records with country information.

### Topical and country impact metrics

Three metrics were used to access topical and country impacts, namely, SCImago Journal Rank (SJR, (*44*)), H-index (the number of articles, *h*, in a journal that have received at least *h* citations), and C-score (citations per document over a two-year period). These metrics between 1999 and 2020 were obtained from the SCImago Scientific Journal Ranking site (https://www.scimagojr.com/journalrank.php). For each plant science record, the SJR, H-index, or C-score of the journal it was published in were used as the impact score for that record. The overall impact score (*I*_m,t.y_) of topic *t* in year *y* using each impact metric *m* (**Data S8**) is defined as:

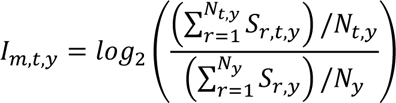

Where *S*_r,t,y_ is the impact score of record *r* in topic *t* published in year *y*, *N*_t,y_ is the number of records in *t* and in *y*, *S*_r,y_ is the impact score of record *r* published in year *y*, and *N*_y_ is the number of records in *y*. The first average was divided by the second to account for year-to-year differences in overall impact scores. The logarithm was applied to facilitate interpretation of the impact score. To determine annual country impact, impact scores were determined in the same way as that for annual topical impact, except that values for different countries were calculated instead of topics (**Data S8**).

To determine topical preference for a country *C*, a 2×2 table was established with the number of records in topic *T* from *C*, the number of records in *T* but not from *C*, the number of non-*T* records from *C*, and the number of non-*T* records not from *C*. A Fisher’s exact test was performed for each *T* and *C* combination. The preference of *T* in *C* was defined as the degree of enrichment calculated as log likelihood ratio of values in the 2×2 table. Topic 5 was excluded because >50% of the countries did not have records for this topic.

### Topical preferences by country

The top 10 countries could be classified into a China-India cluster, an Italy-Spain cluster, and remaining countries (yellow rectangles, **Fig 5E**). The clustering of Italy and Spain is partly due to similar research focusing on allergens (topic 0) and mycotoxins (topic 54) and less emphasis on gene family (topic 23) and stress tolerance (topic 28) studies (**Fig 5F, fig. S9**). There are also substantial differences in topical focus between some countries. For example, plant science records from China tend to be enriched in hyperspectral imaging and modeling (topic 9), gene family studies (topic 23), stress biology (topic 28), and research on new plant compounds associated with herbal medicine (topic 69), but less emphasis on population genetics and evolution (topic 86, **Fig 5F**). In the US, there is a strong focus on insect pest resistance (topic 75), climate, community, and diversity (topic 83), and population genetics and evolution but less focus on new plant compounds. In summary, in addition to revealing how plant science research has evolved over time, topic modeling provides additional insights into differences in research foci among different countries.

